# Rearrangement Scenarios Guided by Chromatin Structure

**DOI:** 10.1101/137323

**Authors:** Sylvain Pulicani, Pijus Simonaitis, Krister M. Swenson

## Abstract

Genome architecture can be drastically modified through a succession of large-scale rearrangements. In the quest to infer accurate ancestral rearrangement scenarios, it is often the case that parsimony principal alone does not impose enough constraints. Thus, the current challenge is to consider more biological information in the inference process. In previous work, we introduced a model for such a task, based on a partition into equivalence classes of the adjacencies between genes. Such a partition is amenable to the representation of spacial constraints as given by Hi-C data. A major open question is the validity of such a model. In this note, we show that the quality of a clustering of the adjacencies based on Hi-C data is directly correlated to the quality of a rearrangement scenario that we compute between *Drosophila melanogaster* and *D. yakuba*.

## 1 Introduction

Genome architecture is modified on a large scale by *rearrangements*. Even fairly distantly related species, such as human and mouse, can share almost all of the same genes yet have drastically different gene orders [4]. These differences are a result of a succession of rearrangements from an ancestral architecture, called a rearrangement *scenario*. In the quest to infer accurate rearrangement scenarios, it is often the case that the parsimony principal alone does not impose enough constraints [3].

When comparing large-scale genome architecture, *syntenic blocks* of similar sequences of genes between a group of species are first inferred using sequence similarity [5]. The *adjacencies* between these blocks are the potential locations for *breakpoints* that rearrangements act on. We have developed methods for choosing more likely rearrangements based on a partition of these adjacencies into disjoint sets (or *colorings*); pairs of adjacencies in the same set are more likely to take part as breakpoints in a rearrangement than pairs between different sets. Thus we weight rearrangements by giving the inter-set rearrangements a weight of 1, and the intra-set rearrangements a weight of 0. The weight of a scenario is the sum of the weights of the constituent moves. With such a model in hand, we posed two optimization problems in [9].

*Problem 1 (*MLS). Minimum Local Scenario

**INPUT:** Sets of pairs *A* and *B* with a coloring of *A*.

**OUTPUT:** A scenario of rearrangements transforming *A* into *B*.

**MEASURE:** The weight of the scenario.

The problem Minimum Local Parsimonious Scenario introduces the constraint that the output is also a parsimonious scenario, *i.e.* a scenario of minimum length.

*Problem 2 (*MLPS). Minimum Local Parsimonious Scenario

**INPUT:** Sets of pairs *A* and *B* with a coloring of *A*.

**OUTPUT:** A *parsimonious* scenario of rearrangements transforming *A* into *B*.

**MEASURE:** The weight of the scenario.

We will use the term MLS and MLPS to denote the weight of the scenario for an optimal solution of the problem.

This note is devoted to showing the biological applicability of these methods. We do this by showing that meaningful partitions of the adjacencies exists. The hypothesis behind our work is that breakpoints that are close in 3D space are more likely to take part in a rearrangement than those which are distant. Evidence supporting this hypothesis has been reported for inter-species rearrangements [10], as well as for somatic rearrangements [12,2].

We test our methods by computing clusters of adjacencies based on Hi-C data for the *Drosophila* species group [8] (see Section 2.1 for a description of Hi-C). The general experimental setup is the following. Genomes of *Drosophila melanogaster* and *Drosophila yakuba* are first partitioned into 64 syntenic blocks. Then the Hi-C data for *melanogaster* is used as a similarity measure on the adjacencies. Clustering around medoids [7] is used to construct colorings based on the similarity function. Then we compute the weights of the MLS and MLPS for the constructed colorings and investigate the relationship between the weights of the clusters and the weights of the scenarios.

We observe a strong correlation between the quality of the clusters and the MLS (*i.e.* the number of moves that are distant according to Hi-C); the better the cluster corresponds to the Hi-C data, the fewer distant moves must be done in a rearrangement scenario. We also study other features of the data that have implications on the computability of the MLS and more general weight functions.

## 2 Methods

### 2.1 Hi-C Data

The Hi-C experiment provides a rough estimate of how many times a pair of genomic loci are in close proximity [6]. Roughly speaking, formaldehyde is introduces to a population of cells so that parts of the genomes that are in spacial proximity are linked together. Each side of the link contains segments of DNA that are then sequenced. The sequences are mapped to a reference genome so that the pair of spacially proximal loci are determined. Thus, each cell produces a set of pairs of loci. With a large population of cells, we have a distribution of pairs that are finally corrected for experimental biases.

Due to the nature of contact probability, which decreases dramatically with respect to chromosomal distance (it roughly follows a power law), we applied the normalization done by Lieberman-Aiden *et al.* (see the appendix of [6]) to the matrices obtained from [8]. This allows the long rearrangements (in the genetic coordinate sense) to have the same importance as the short ones. Specifically, a normalized intrachromosomal heatmap entry *INTRA*_*ij*_ gets the value

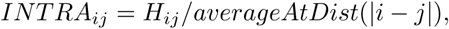

where *averageAtDist*(*d*) is the expected value of an entry *d* off of the diagonal of any intrachromosomal matrix. A normalized interchromosomal heatmap entry *INTER*_*ij*_ gets the value

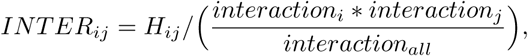

where *interaction*_*x*_ is the sum of all interaction for position *x*, and *interaction*_*all*_ is the total sum of all entries in all matrices.

### 2.2 Clustering

Clustering around medoids [7] was chosen for its simplicity and speed. We use the Hi-C data as a similarity measure for the clustering; the more Hi-C contacts there are between two adjacencies, the more likely we are to have the two in the same cluster. A *medoid* of a cluster is an element that maximizes the sum of the similarities to the rest of a cluster. This sum is the cluster’s *weight*, and when summed over all the clusters it provides us with a *weight function* for a clustering. An important property of this clustering method is that it provides many local optima that we can compare to MLS. We use three clustering algorithms: K-MEDOIDS, RANDOM and MIXED that generate *k* clusters for a chosen *k* which in our case range from 1 to 70.

The K-MEDOIDS algorithm starts with randomly initialised centroids. The rest of the elements are then associated to the centroids that are most similar to them. The medoids of the obtained clusters are then computed and they become the new centroids around which the elements will be clustered. We continue this procedure until the weight of a clustering stops increasing.

The RANDOM algorithm partitions the elements at random into *k* non-empty clusters. On our data we observe that the weights of the random clusters are always smaller than those provided by the clustering around medoids and in order to bridge the gap between the obtained weights we mix K-MEDOIDS and random algorithms to obtain a mixed algorithm that initialises the centroids randomly and then chooses at random how many of the resulting elements will get assigned to the clusters based on the similarity function, and how many of them will be assigned at random (without performing further iterations).

### 2.3 Creation of Syntenic Blocks

We computed syntenic blocks in two steps. First, we took the orthologs for *D. melanogaster* and *D. yakuba*. This was done using the OMA groups database [1]. We removed each gene that overlaps or intersects another, along with its ortholog in the other species. Then, the blocks were constructed using the Orthocluster tool [11]. The basic idea is to aggregate orthologs to make the biggest possible blocks without breaking certain constraints that define the synteny; the constraints are the maximal and minimal number of genes per blocks, the absolute gene order between the genomes, the genes strandedness, the quantity of non-ortholog genes and the possibility to make nested blocks. We forbid the creation of nested blocks as we wanted a 1-1 block mapping. All other parameters were default. Orthocluster outputs clusters of genes. In order to get back syntenic blocks, we take the smallest gene position in a cluster as the start position of the block from this cluster, and the biggest position as the end for that block. We then check back the blocks for the presence of intersection and overlap. We did not find any.

## 3 Results

### 3.1 MLS and the weight of a coloring

Figure 1 presents 200 independent clusterings of the adjacencies of *Drosophila melanogaster* into *k* = 15 clusters. The blue half of them is generated using random, the rest using the K-MEDOIDS algorithm. There is a clear separation on both axes between these two sets of clusterings. The MLS and cluster weight is always significantly better on a K-MEDOIDS clustering than a random clustering. Figure 7 to Figure 8 show such plots for other values of *k*.

**Fig. 1.**
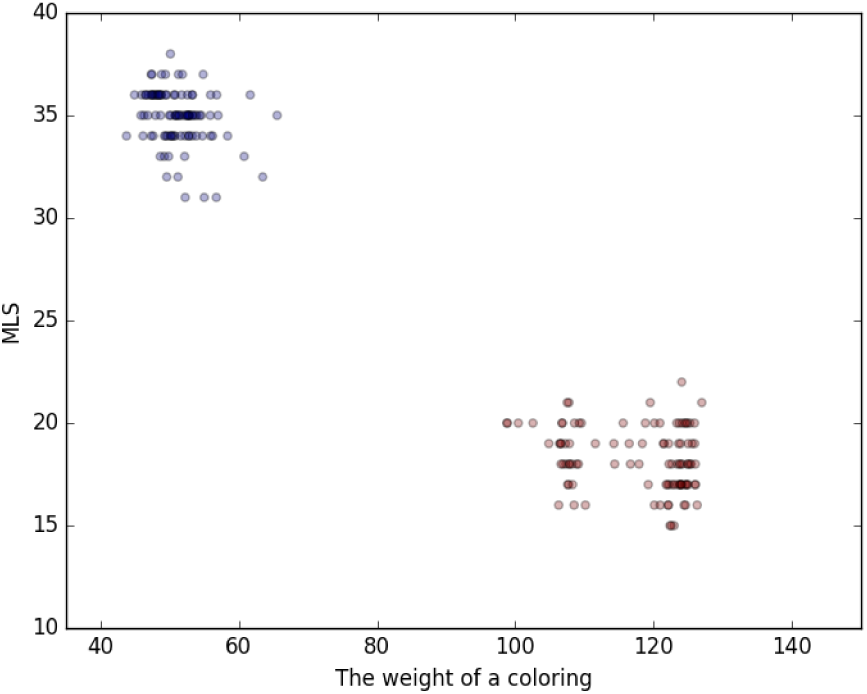
For k = 15 on *melanogaster* Hi-C data, random clusters are in blue while K-MEDOIDS clusters are in red.

The MIXED algorithm was introduced in order to bridge the gap between the weights of the colorings provided by RANDOM and K-MEDOIDS. We note a significant inverse correlation between the weights of the colorings and MLS. Figure 2 depicts 100 independent clusterings for the adjacencies of *Drosophila melanogaster* obtained using the mixed algorithm. In this plot a color of a point indicates how many of the adjacencies in that particular clustering got assigned to the clusters at random during a run of mixed. Blue shows that a clustering is mostly random and red, on the other hand, means that most of the adjacencies got assigned to the centroids based on the similarity function. In this example Pearson’s correlation *r* is found to be -0.87 with a 2-tailed p-value of 1.0-31. Figure 9 to Figure 12 present experiments for different values of *k* showing that the correlation first increases with *k* before slowly decreasing.

**Fig. 2.**
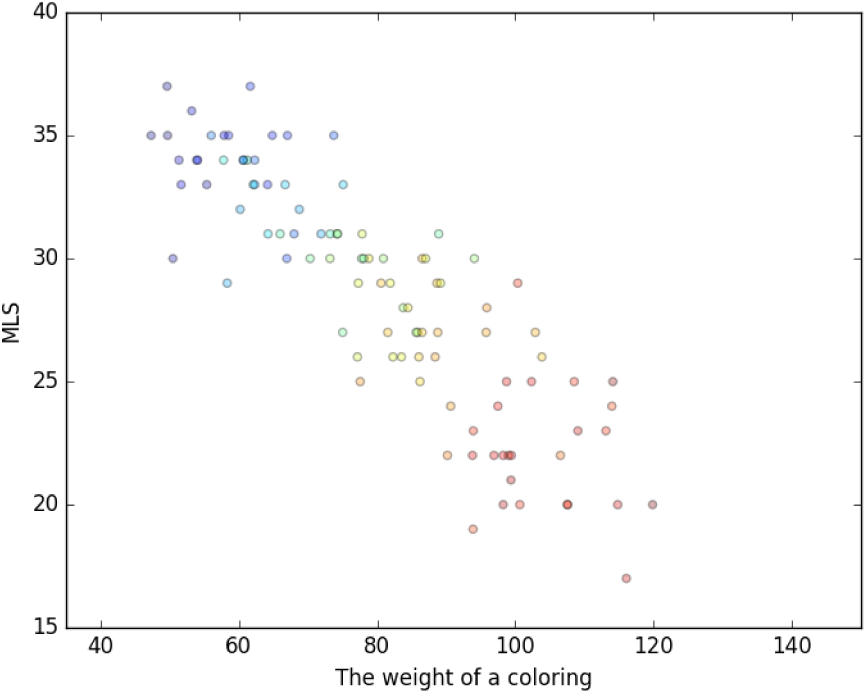
For k = 15 on *melanogaster*, MIXED clusters with varying amounts of randomly assigned adjacencies. As more randomization is introduced to the clustering the color of the points moves from red to yellow to green to blue. For this data the Pearson’s correlation r = -0.87, and the p-value = 1.0-31.

### 3.2 Computational complexity

In general, finding MLS is computationally costly, as we have proven it to be NP-hard. In other words, we cannot expect to be able to compute a scenario that minimizes the number of non-local moves on any pair of genomes. We established an exact algorithm, however, that runs efficiently if a certain parameter called the number of *simple cycles* is “small enough”. We expect “small enough” to roughly be in the hundreds of thousands. Between *melanogaster* and *yakuba*, we find that the number of simple cycles is always very small using the clusters computed by the K-MEDOIDS algorithm. In particular, we ran 100 instances of K-MEDOIDS using Hi-C data from *melanogaster* for every *k* ranging from 2 to 70, and computed the number of simple cycles for those clusterings. The values for all 6900 runs are presented in Figure 3. The average number of simple cycles is 16.9 for *k* = 42, the maximum that we observed. The average number of simple cycles over all *k* is 8.5. We conjecture that this value would also be low enough for human Hi-C data on scenarios between human and mouse.

**Fig. 3.**
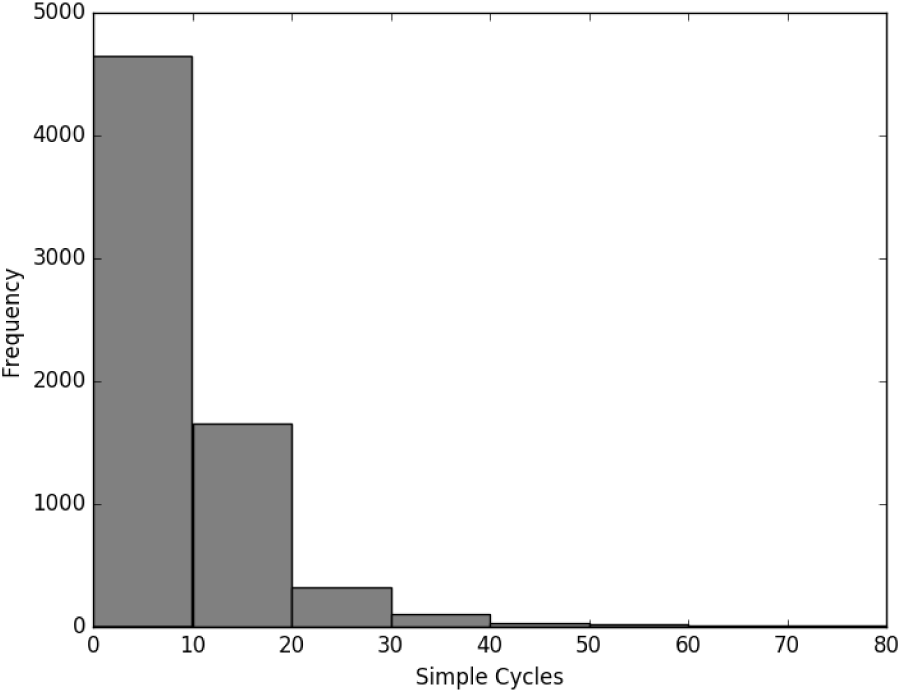
K-MEDOIDS clusters for *melanogaster*, average = 8.5, standard deviation = 8.4, highest average of 16.9 for *k* = 42

### 3.3 MLPS – MLS and the weight of a coloring

We study the value of the difference MLPS *-* MLS due to its potential role in computing a more general weight function. When this difference is low, the number of non-parsimonious rearrangements is few, and a more general weight function on non-local moves is easier to compute and approximate.

A significant correlation between the difference MLPS *-* MLS and the weights of the colorings is also found, as depicted in Figure 4 and Figure 5 for *k* = 10, and in Figures 13 to 16 for other values of *k*.

**Fig. 4.**
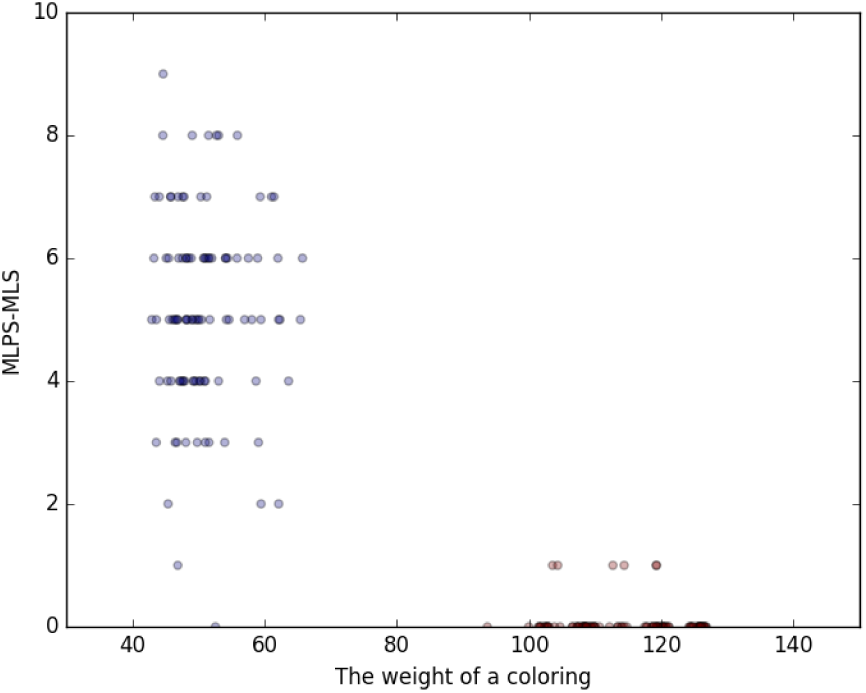
k = 10, RANDOM and K-MEDOIDS clusters for *melanogaster*

**Fig. 5.**
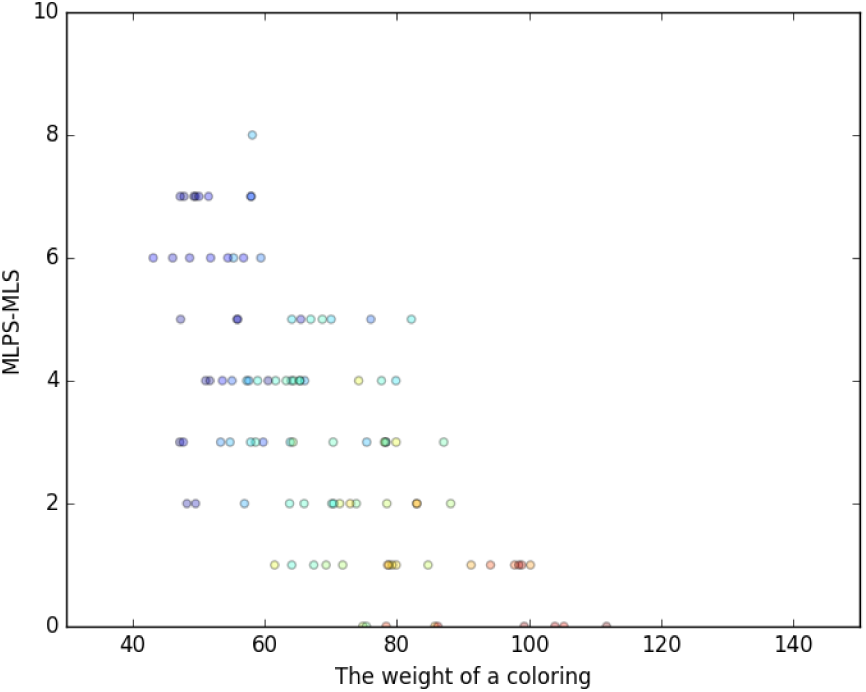
k = 10, MIXED clusters for *melanogaster*, r = -0.69, p-value = 2.0-15 Further, for the clusterings provided by K-MEDOIDS this value of MLPS *-* MLS is low in general, as displayed in Figure 6. As for Figure 3 we did 100 runs of K-MEDOIDS for every *k*, and found the average for MLPS *-* MLS to be highest at *k* = 24 with a value 0.29. The average over all *k* was 0.12.

**Fig. 6.**
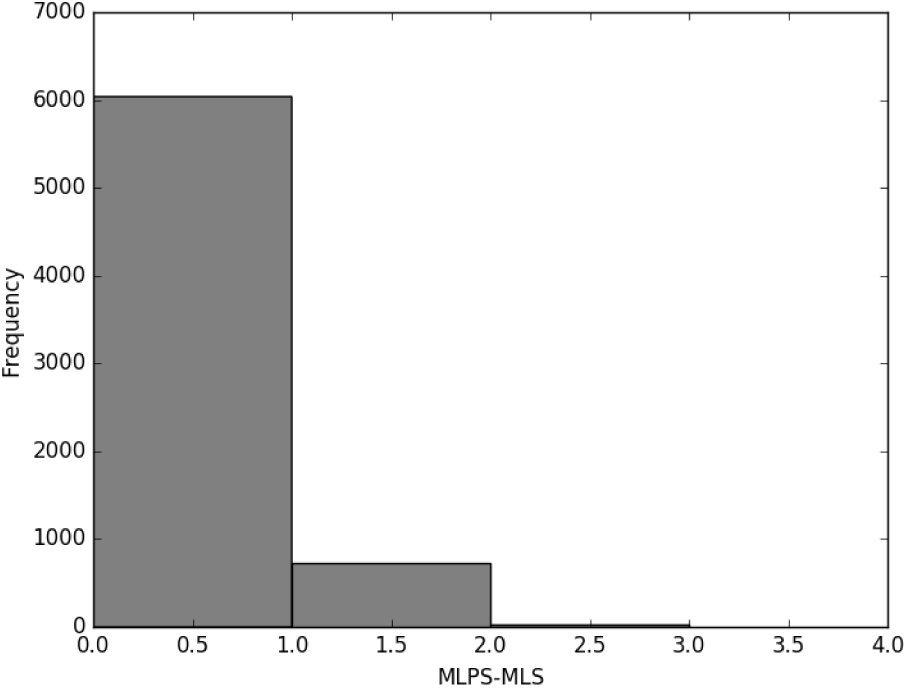
K-MEDOIDS clusters for *melanogaster*, average = 0.12, standard deviation = 0.34, highest average of 0.29 for *k* = 24

Figure 17 presents a similar histogram for the runs of RANDOM for *melanogaster*. As expected, RANDOM clusterings produce larger numbers of simple cycles.

**Fig. 7.**
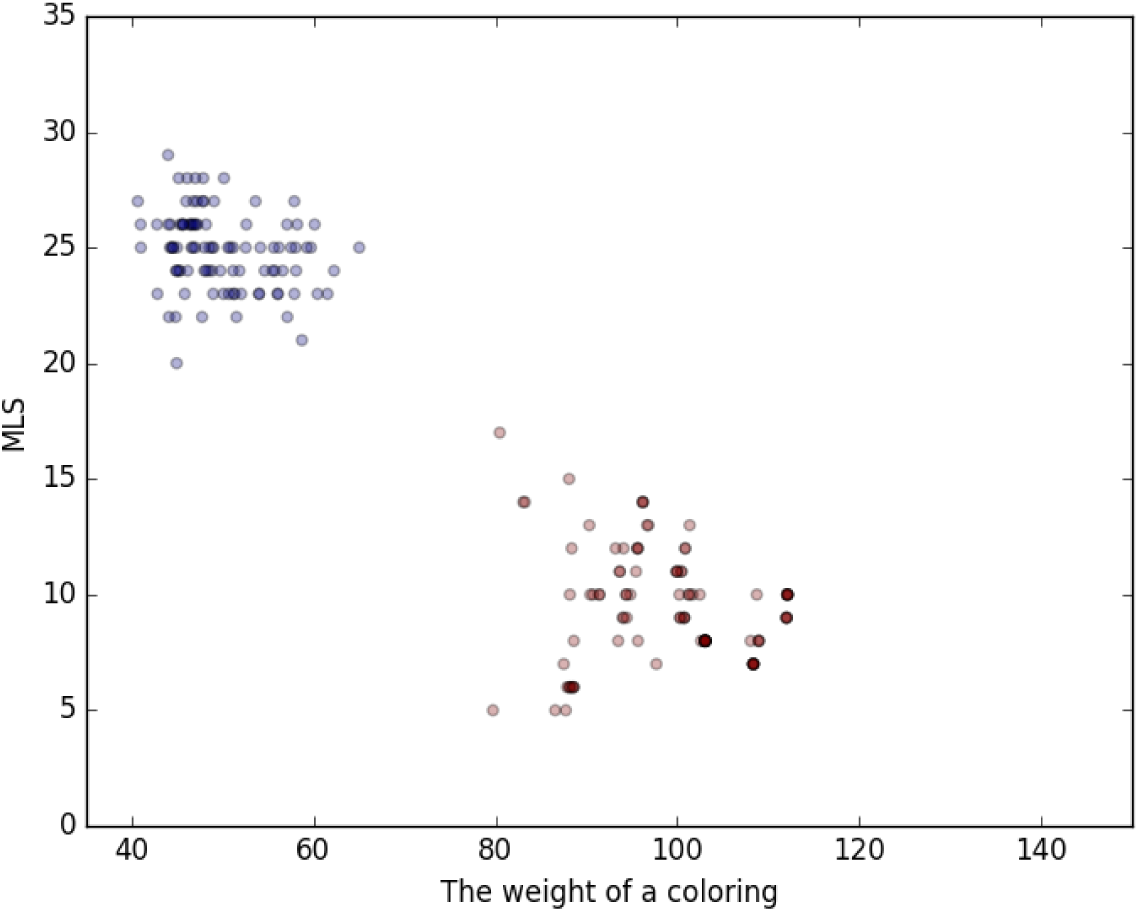
k = 5, RANDOM and K-MEDOIDS clusters for *melanogaster*

**Fig. 8.**
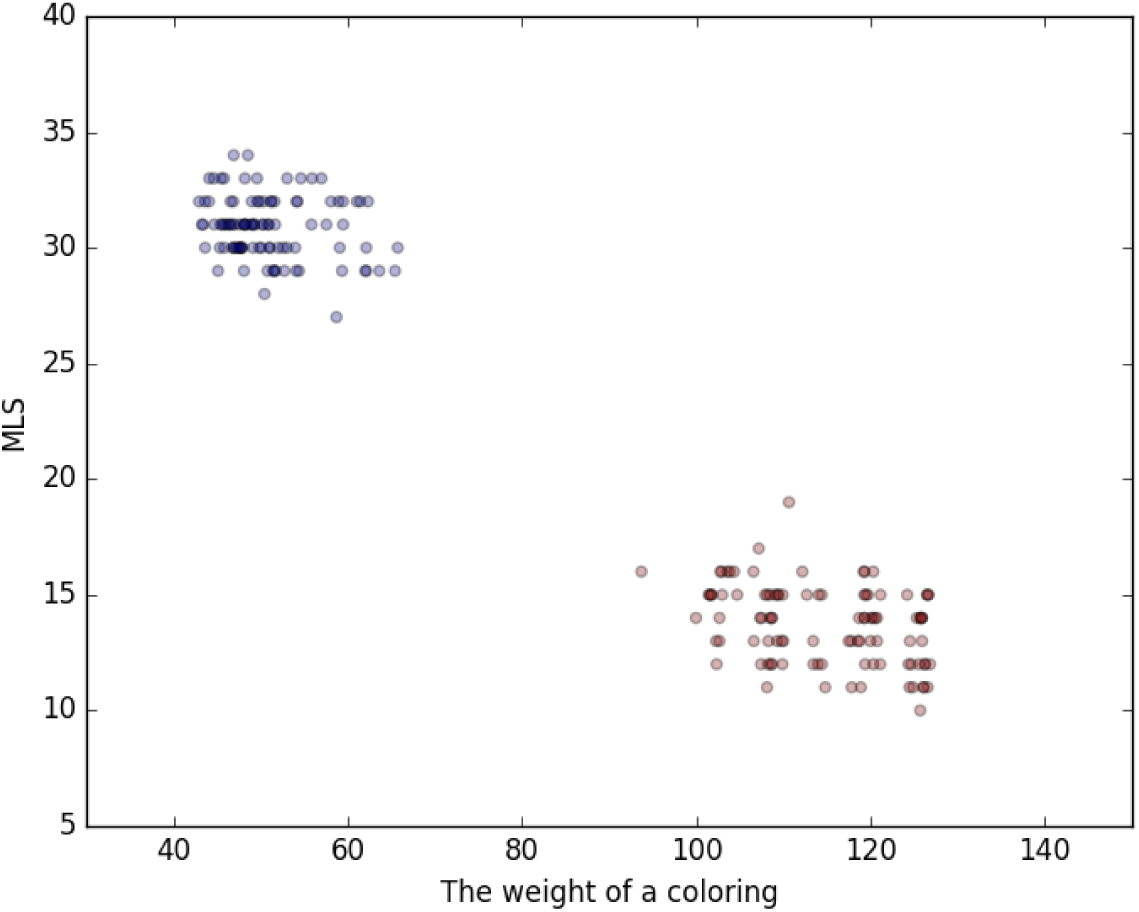
k = 10, RANDOM and K-MEDOIDS clusters for *melanogaster*

**Fig. 9.**
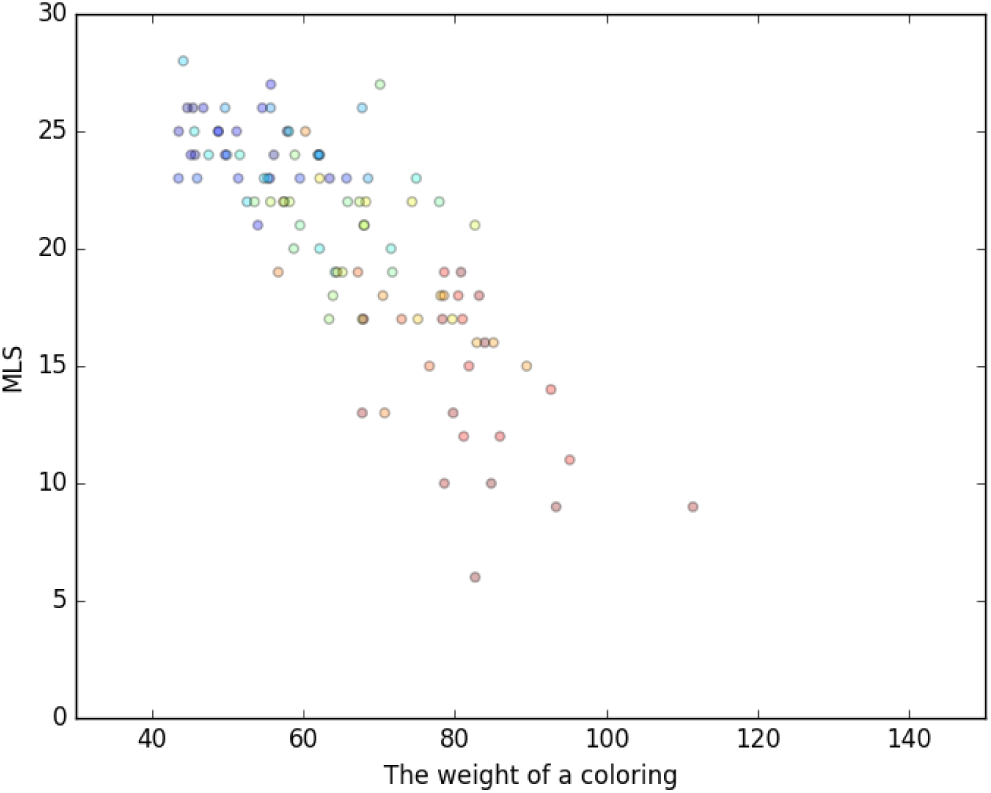
k = 5, MIXED clusters for *melanogaster*, r = -0.80, p-value = 1.3-23

**Fig. 10.**
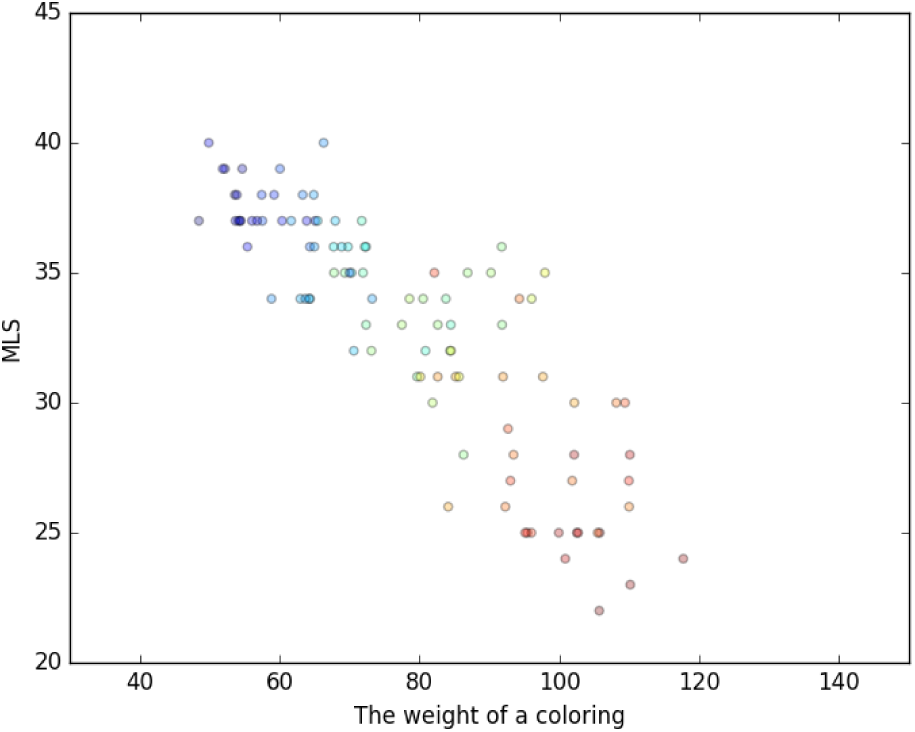
k = 20, MIXED clusters for *melanogaster*, r = -0.85, p-value = 1.0-29

**Fig. 11.**
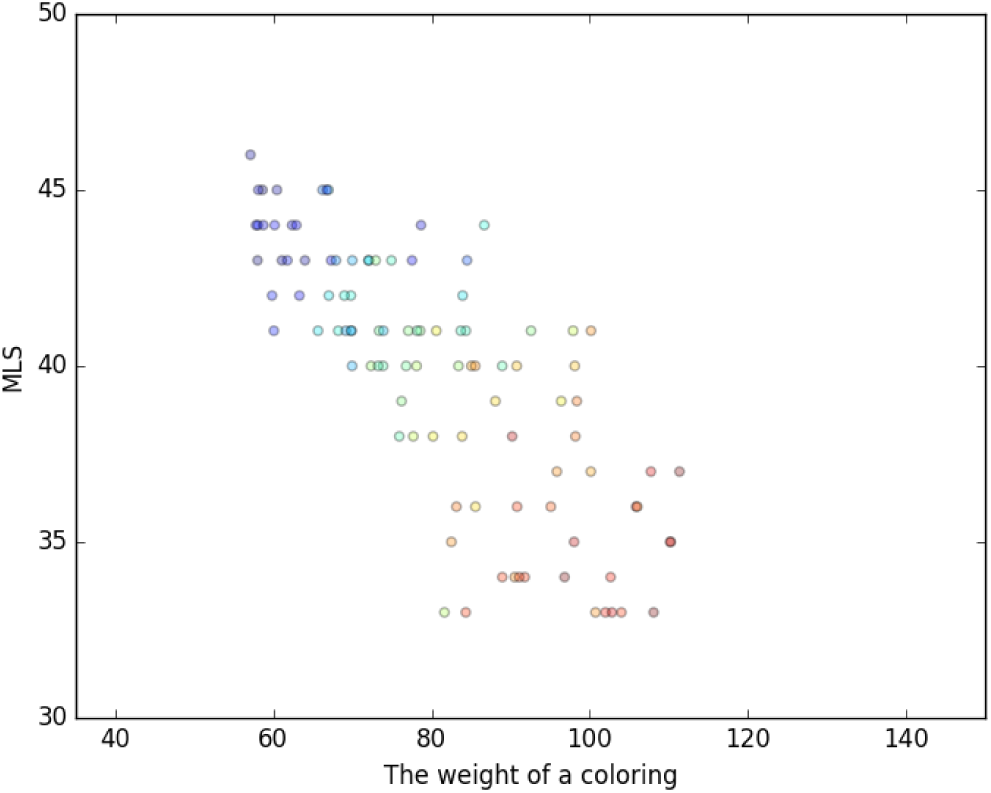
k = 35, MIXED clusters for *melanogaster*, r = -0.77, p-value = 3.4-21

**Fig. 12.**
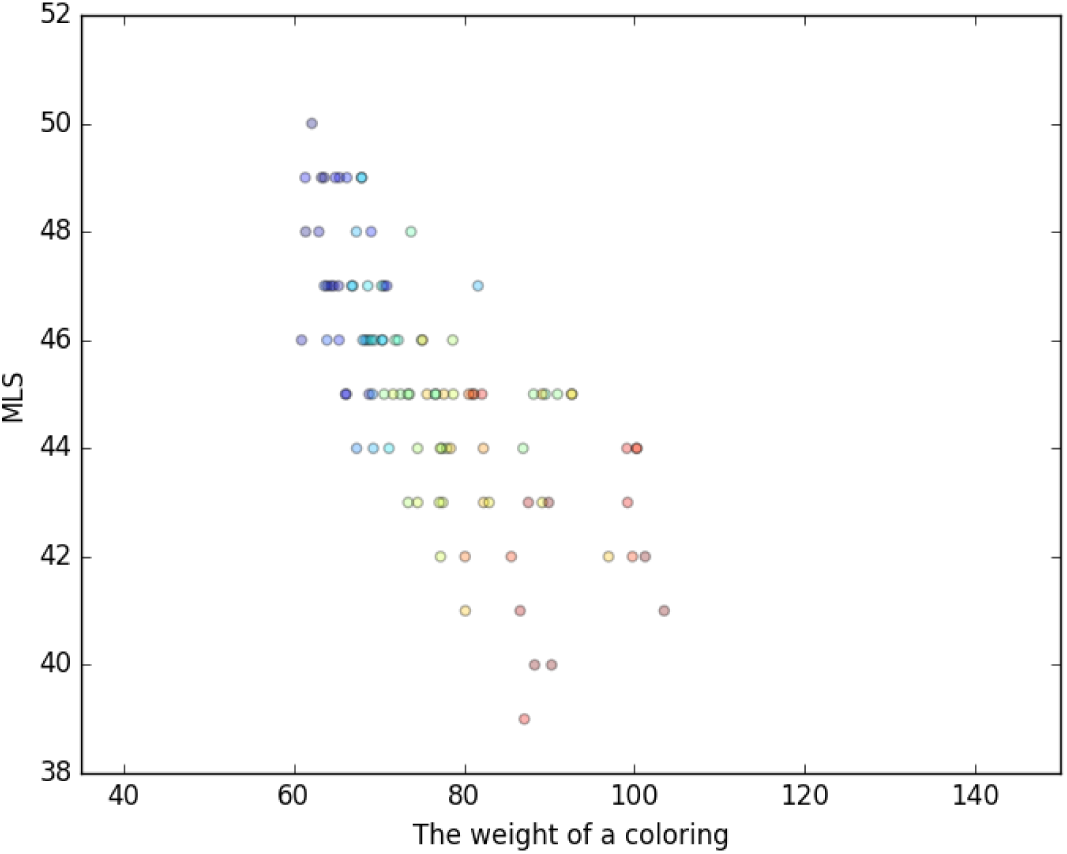
k = 50, mixed clusters for *melanogaster*, r = -0.69, p-value = 5.7-16

**Fig. 13.**
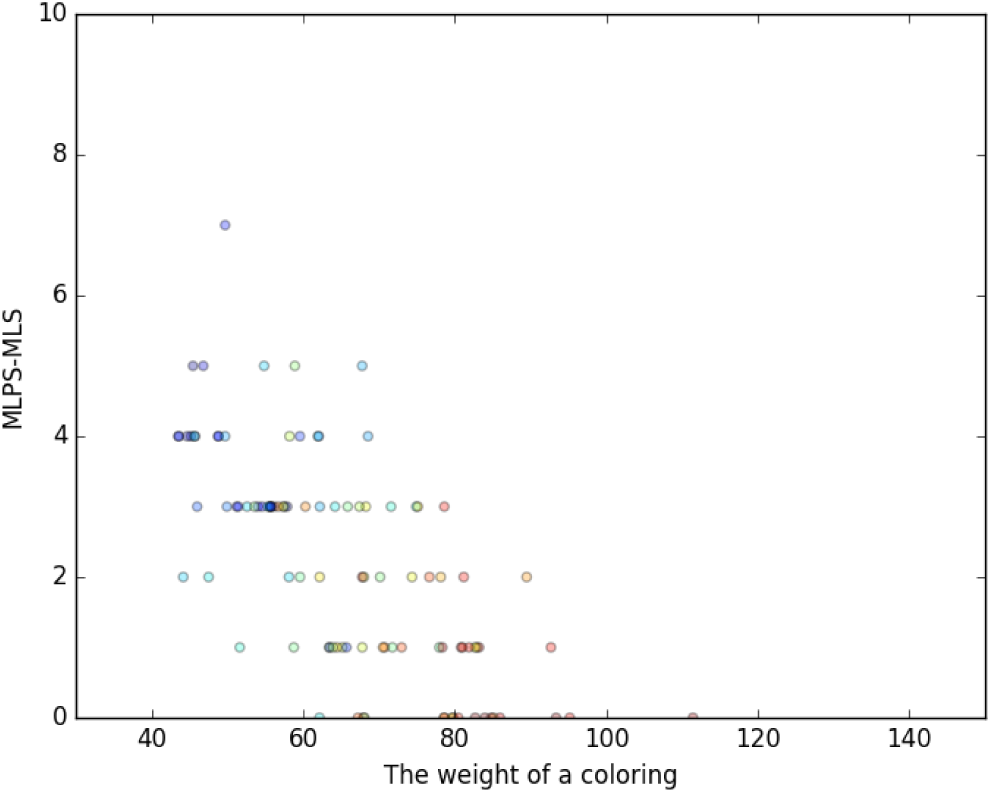
k = 5, MIXED clusters for *melanogaster*, r = -0.69, p-value = 1.8-15

**Fig. 14.**
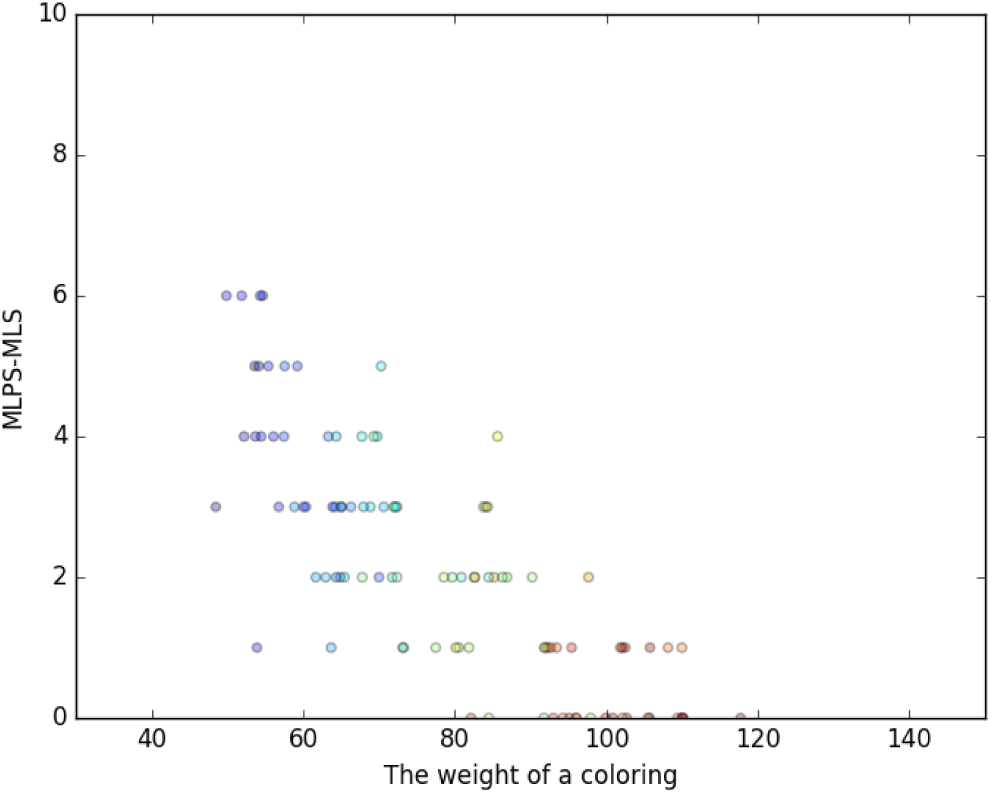
k = 20, MIXED clusters for *melanogaster*, r = -0.79, p-value = 3.1-23

**Fig. 15.**
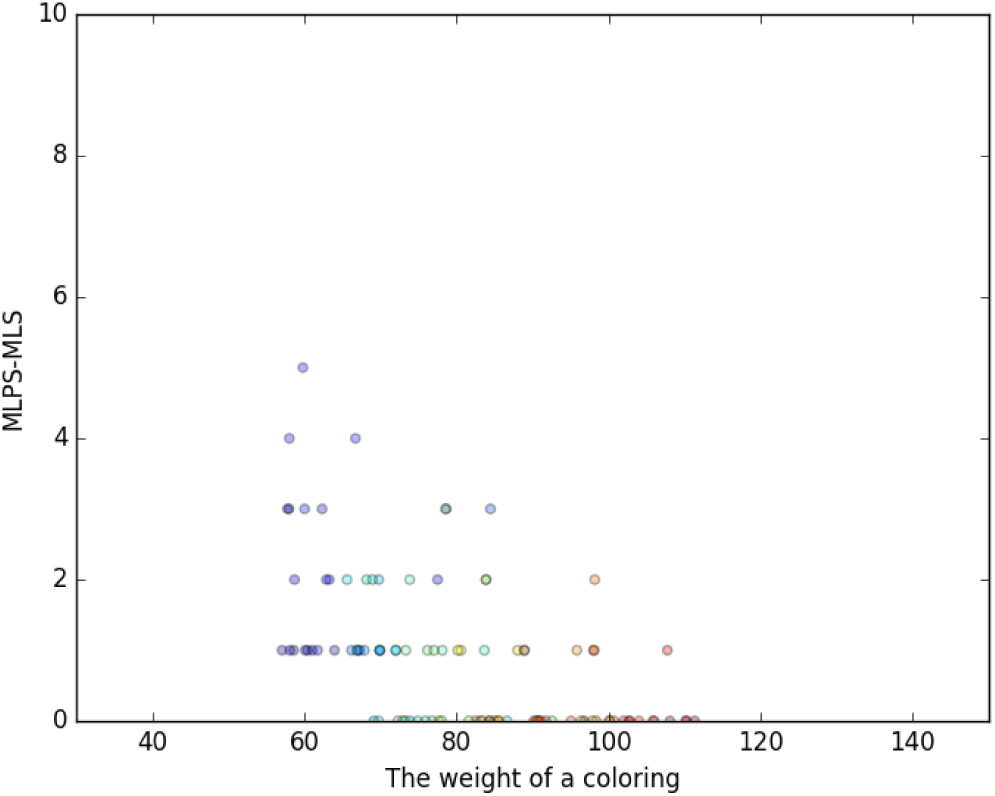
k = 35, mixed clusters for *melanogaster*, r = -0.54, p-value = 5.0-09

**Fig. 16.**
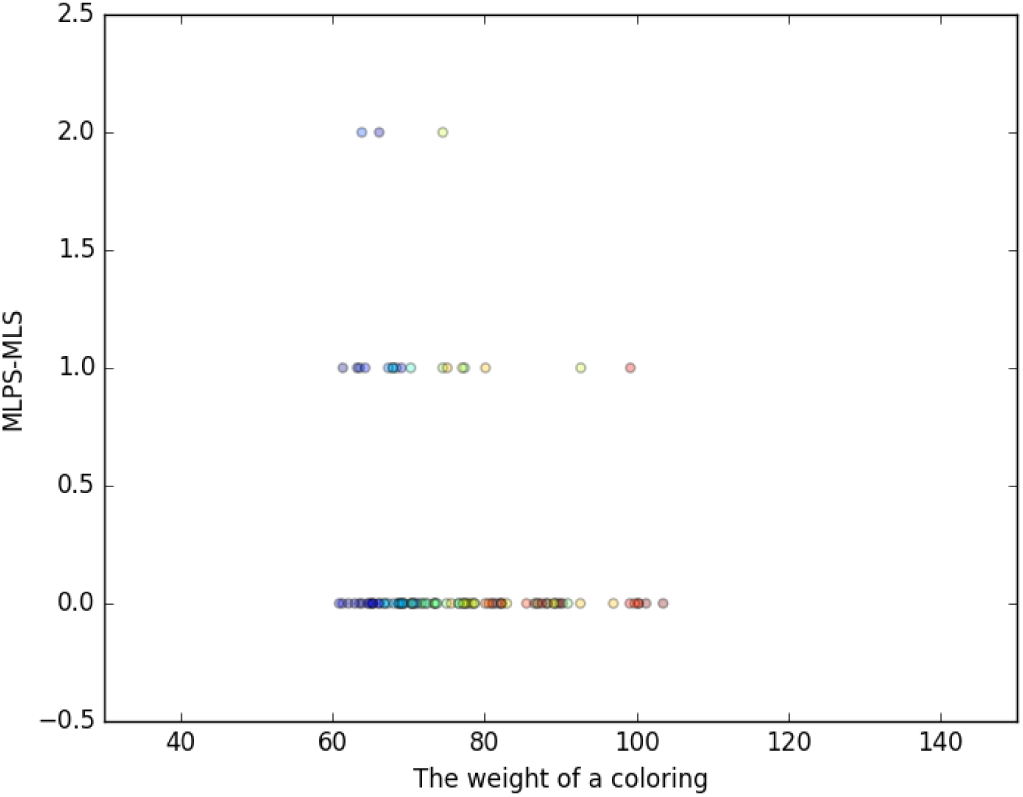
k = 50, mixed clusters for *melanogaster*, r = -0.21, p-value = 0.04

For every *k* ranging from 2 to 70 we ran an instance of random and computed the number of the simple cycles, as well as the difference MLPS – MLS. The computed histograms are presented in Figures 17 and 18.

**Fig. 17.**
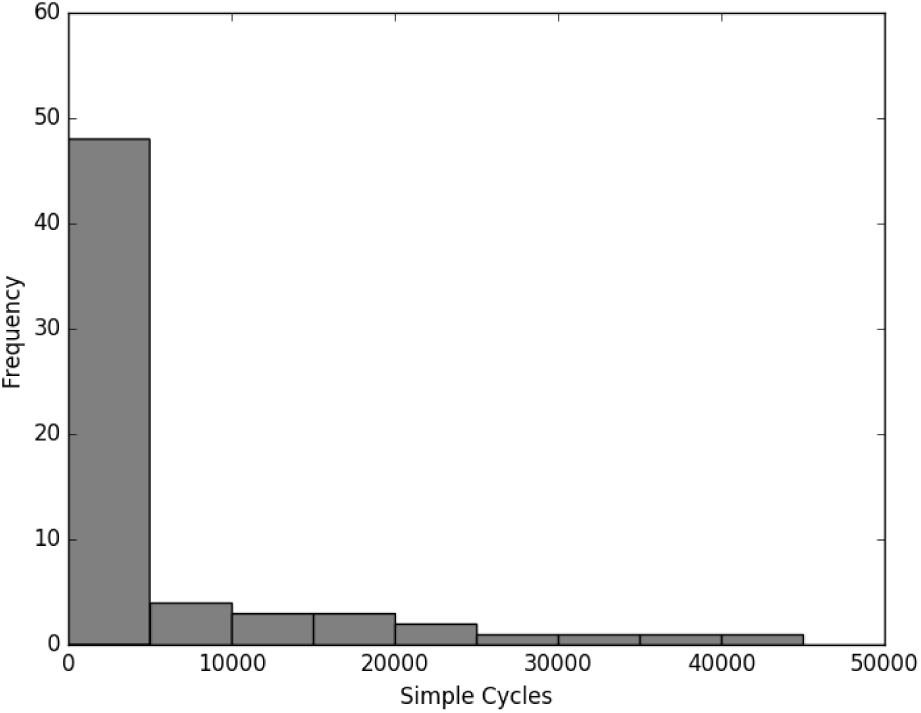
RANDOM clusters for *melanogaster*. The average number of simple cycles is 9164, standard deviation is 18,227. Four values higher than 50,000 were detected with a maximum of 86319.

**Fig. 18.**
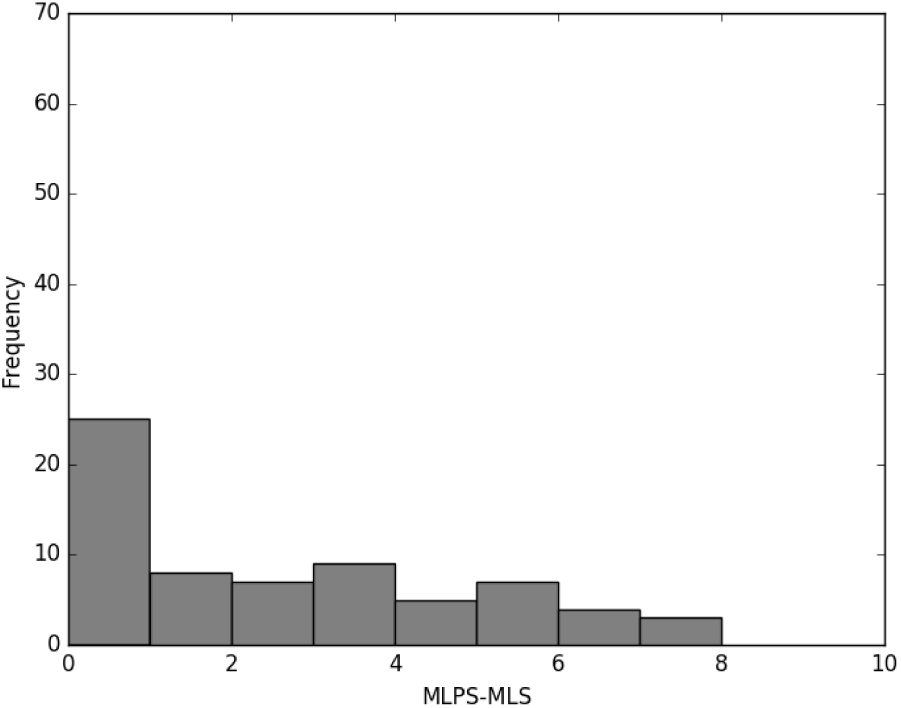
RANDOM clusters for *melanogaster*, average = 2.19, standard deviation = 2.2, and the highest value is 7.

A similar histogram for the runs of RANDOM for *melanogaster* can be found in Figure 18

## 4 Conclusions and Future Work

We have presented a strong correlation between the quality of a clustering of genomic adjacencies, and the number of DCJ rearrangements that must be done between these clusters. This shows that partitions on genome adjacencies can be constructed using Hi-C data which inform rearrangement scenarios through the use of the Minimum Local Scenario and Minimum Local Parsimonious Scenario problems.

For now, we have used a simple weight function and a simple K-MEDOIDS clustering [7]. In the future better clusterings can be computed to suit our needs for high within-cluster similarity and a dissimilarity between the clusters. Other clustering techniques, such as hierarchical clustering, might also be of interest.

From a practical perspective, results showing that the difference MLPS – MLS is always low give hope that a more general weight function could be developed.

## 5 Acknowledgements

Part of this work was funded by NUMEV grant AAP 2014-2-028 and EPIGEN-MED grant ANR-10-LABX-12-01.

## References

1. Adrian M Altenhoff, Nives Škunca, Natasha Glover, Clément-Marie Train, Anna Sueki, Ivana Pili΂ota, Kevin Gori, Bartlomiej Tomiczek, Steven Müller, Henning Redestig, et al. The oma orthology database in 2015: function predictions, better plant support, synteny view and other improvements. Nucleic acids research, page gku1158, 2014.

2. Peter J Campbell, Philip J Stephens, Erin D Pleasance, Sarah O’Meara, Heng Li, Thomas Santarius, Lucy A Stebbings, Catherine Leroy, Sarah Edkins, Claire Hardy, et al. Identification of somatically acquired rearrangements in cancer using genome-wide massively parallel paired-end sequencing. Nature genetics, 40(6):722–729, 2008.

3. Cedric Chauve, Haris Gavranovic, Aida Ouangraoua, and Eric Tannier. Yeast ancestral genome reconstructions: the possibilities of computational methods ii. Journal of Computational Biology, 17(9):1097–1112, 2010.

4. Asif T Chinwalla, Lisa L Cook, Kimberly D Delehaunty, Ginger A Fewell, Lucinda A Fulton, Robert S Fulton, Tina A Graves, LaDeana W Hillier, Elaine R Mardis, John D McPherson, et al. Initial sequencing and comparative analysis of the mouse genome. Nature, 420(6915):520–562, 2002.

5. Cristina G Ghiurcuta and Bernard ME Moret. Evaluating synteny for improved comparative studies. Bioinformatics, 30(12):i9–i18, 2014.

6. Erez Lieberman-Aiden, Nynke L. van Berkum, Louise Williams, Maxim Imakaev, Tobias Ragoczy, Agnes Telling, Ido Amit, Bryan R. Lajoie, Peter J. Sabo, Michael O. Dorschner, Richard Sandstrom, Bradley Bernstein, M. A. Bender, Mark Groudine, Andreas Gnirke, John Stamatoyannopoulos, Leonid A. Mirny, Eric S. Lander, and Job Dekker. Comprehensive mapping of long-range interactions reveals folding principles of the human genome. Science, 326(5950):289–293, Oct 2009.

7. Hae-Sang Park and Chi-Hyuck Jun. A simple and fast algorithm for k-medoids clustering. Expert Systems with Applications, 36(2, Part 2):3336 – 3341, 2009.

8. Tom Sexton, Eitan Yaffe, Ephraim Kenigsberg, Frédéric Bantignies, Benjamin Leblanc, Michael Hoichman, Hugues Parrinello, Amos Tanay, and Giacomo Cavalli. Three-dimensional folding and functional organization principles of the drosophila genome. Cell, 148(3):458–472, 2012.

9. Krister M. Swenson, Pijus Simonaitis, and Mathieu Blanchette. Models and algorithms for genome rearrangement with positional constraints. Algorithms for Molecular Biology, 11(1):13, 2016.

10. Amélie S Véron, Claire Lemaitre, Christian Gautier, Vincent Lacroix, and Marie-France Sagot. Close 3d proximity of evolutionary breakpoints argues for the notion of spatial synteny. BMC genomics, 12(1):303, 2011.

11. Xinghuo Zeng, Matthew J Nesbitt, Jian Pei, Ke Wang, Ismael A Vergara, and Nansheng Chen. Orthocluster: a new tool for mining synteny blocks and applications in comparative genomics. In Proceedings of the 11th international conference on Extending database technology: Advances in database technology, pages 656–667. ACM, 2008.

12. Yu Zhang, Rachel Patton McCord, Yu-Jui Ho, Bryan R Lajoie, Dominic G Hildebrand, Aline C Simon, Michael S Becker, Frederick W Alt, and Job Dekker. Spatial organization of the mouse genome and its role in recurrent chromosomal translocations. Cell, 148(5):908–921, 2012.

